# Identification of QTLs linked to partial resistance to foot and root rot caused by *Fusarium avenaceum* and *Fusarium oxysporum* in faba bean (*Vicia faba*)

**DOI:** 10.1101/2025.02.21.639454

**Authors:** Anne Webb, Jane E. Thomas, Huw Davis, Simon McAdam, Deepti Angra, Donal M. O’Sullivan, Krystyna Gostkiewicz, Martina Capozzi, Thomas A. Wood

## Abstract

Foot and root rot, caused by a complex of soil-borne fungal and oomycete pathogens, including several *Fusarium* spp., can cause serious yield losses in faba bean. Current control strategies rely largely on agronomic practices and limited varietal resistance. Identifying novel sources of genetic resistance is of great potential value for breeding elite varieties with improved foot rot resistance. A partially resistant *Vicia faba* accession ‘ig124213’ (NV490) was crossed with a moderately susceptible ‘ig124301’ (NV512) and a mapping population comprising 198 F_3_ families was developed. This was screened for resistance to a mixture of seven UK field isolates of *Fusarium avenaceum* and *Fusarium oxysporum* under glasshouse conditions. Several F_3_ families with moderate to high levels of resistance to both *Fusarium* species were identified, and a high-density linkage map of the *V. faba* genome including 6755 SNP-markers in seven linkage groups (LG) was generated using the ‘Vfaba_v2’ Axiom SNP array. Quantitative trait loci associated with improved resistance to *Fusarium* foot and root rot were identified, including one major QTL on LG4, corresponding to the chromosome 4 of *V. faba*.

**Key message:** The first QTL associated with partial resistance to *Fusarium* foot and root rot has been identified using a biparental mapping population of *Vicia faba*.

## Introduction

Foot and root rot in faba bean (*Vicia faba* L.) is a soil-borne and possibly also seed-borne (McGee and Kellock 1974; Pande et al. 2007) disease caused by the pathogen complex comprising several fungal and oomycete species. This is thought to include *Rhizoctonia solani, Didymella pinodella*, and *Aphanomyces euteiches*, as well as several *Fusarium* species, e. g. *F. solani, F. oxysporum, F. avenaceum, F. clavum, F. culmorum, F. equiseti, F. redolens* and *F. vanettenii* (Clarkson 1978; Lamari and Bernier 1985; Afshari et al. 2020; Dugassa et al. 2021; Šišić et al. 2022, Attar et al. 2024). Multiple *Fusarium* species have been found to be causing foot and root rot in the same field and the species complex composition can vary with soil type, management practices, geographic location, humidity and temperature (Esmaeili Taheri et al. 2017; Šišić et al. 2022; Yu et al. 2023), highlighting the importance of broad-spectrum resistance as a valuable breeding target. Disease symptoms can appear at any growth stage of the crop. Affected seedlings either fail to emerge after germination or show stunted growth with yellowing of foliage, progressing to dry papery necrosis and death before flowering. In adult plants symptoms may begin with initially reversible wilting in warm, dry conditions, followed by chlorosis and necrosis appearing in lower leaves with brown or black stem lesions becoming apparent at the base of the plant, progressing to premature senescence and death, leading to a large reduction or even a total loss of yield (Yu et al. 2023).

Underground symptoms caused by *Fusarium* spp. in faba bean include brown or black discolouration of the hypocotyl and roots, with white, peach or orange coloured mycelium appearing on dead stem tissue. Within fields, the disease appears in gradually enlarging patches ranging in size from few scattered individual plants to large bare patches. Seed treatment with fungicides (Chang et al. 2014), intercropping and nitrogen supplementation (Lv et al. 2021) and biocontrol agents (El-Mougy and Abdel-Kader 2008; Haddoudi et al. 2021) have shown promise in experimental settings. Against the background of reduced availability of active ingredients, alternative means of control must be identified including introduction of broad-spectrum resistance into elite varieties. Several sources of partial resistance to *Fusarium* foot and root rot are known in *V. faba* (Mahmoud and Abd El-Fatah 2020), but no resistance quantitative trait loci (QTLs) have yet been identified. Several QTLs have been identified in other legumes, conferring either broad, race non-specific resistance or race-specific resistance to *Fusarium* spp. have been identified, including in chickpea (*Cicer arietinum*) (Jendoubi et al. 2016; Garg et al. 2018), lentil (*Lens culinaris*) (Hamwieh et al. 2005), pea (*Pisum sativum*) (Grajal-Martìn and Muehlbauer 2002; Mc Phee et al. 2012; Coyne et al. 2015, 2019; Wu et al. 2022) and common bean (*Phaseolus vulgaris*) (Nakedde et al. 2016; Wang et al. 2018; Paulino et al. 2021).

The generation of high-density genetic linkage maps in several legumes, such as faba bean, field pea, lentil, chickpea and common bean (Duarte et al. 2014; Kaur et al. 2014; Paulino et al. 2021; Heineck et al. 2022; Alsamman et al. 2024), has greatly advanced the identification of QTLs and markers associated with desirable agronomic traits. Moreover, genomes of two *V. faba* cultivars, ‘Hedin/2’ and ‘Tiffany’, have been published (Jayakodi et al. 2023) which will greatly assist the identification and characterization of candidate genes controlling desirable phenotypes such as disease resistance, in addition to exploring synteny with closely related, more extensively studied legume species.

Isolates of the two *Fusarium* species used in this study, *F. avenaceum* and *F. oxysporum*, have been sourced from symptomatic faba bean plants grown in Cambridgeshire, UK. An F_3_ population derived from a cross between two inbred faba bean accessions originating from Tajikistan, a partially resistant line NV490-3 (‘ig124213’) and a more susceptible line NV512-1 (‘ig124301’), was evaluated for resistance to a mixture of seven isolates of *F. avenaceum* and *F. oxysporum* in two replicated glasshouse-based experiments. 184 individuals from the F_2_ generation were genotyped on the Vfaba_v2 Axiom SNP array (O’Sullivan et al. 2019; Khazaei et al. 2021). A high-density linkage map, comprising 6577 markers in seven linkage groups representing the six chromosomes of *V. faba*, was generated, allowing the identification of QTLs linked to partial resistance to *Fusarium* foot rot for the first time.

## Materials & Methods

### Population development

Two inbred *V. faba* accessions originating from Tajikistan, NV490-3 (ig124213, partially resistant) and NV512-1 (ig124301, susceptible), obtained from ICARDA, Aleppo, Syria, were crossed and advanced to F_2_, yielding a progeny of 4376 F_2_ seeds. Of these, 184 individuals were grown to F_3_ for phenotyping.

### *Fusarium* isolates

Seven isolates of *Fusarium* spp. were obtained from lower stem tissue of symptomatic faba bean plants (suppl. table S1). Stem sections were surface sterilised with 10 % sodium hypochlorite solution for 15 min, plated on potato dextrose agar (PDA) and maintained at room temperature. Individual isolates were obtained from hyphal tips. For species identification, DNA was extracted from freeze dried mycelium using the Macherey-Nagel Nucleospin Plant II kit (Macherey-Nagel, Düren, Germany) with the extraction protocol modified to include an additional chloroform purification step. After tissue lysis, 250 µl chloroform:isoamyl alcohol (4:1) were added to the lysate, followed by 10 s of vortexing and centrifugation at 20,000 x *g* for 15 min. A partial sequence of *translation elongation factor 1 alpha* (*TEF1α*) gene from each isolate was amplified following the protocol of Taylor et al. (2016) and sequenced from both directions at Azenta Life Sciences (Takeley, UK). Sequences were searched against the FUSARIUM ID v. 3.0 database (Geiser et al. 2004; Torres-Cruz et al. 2022) and the NCBI Blastn nucleotide database (Altschul et al. 1990; Camacho et al. 2009) for species identification.

### Genotyping and construction of *V. faba* linkage map

DNA was extracted from young leaves of 184 F_2_ individuals and the parental lines NV490-3 and NV512-1 using the method described in Doyle and Doyle (1987). 184 samples and two replicates of each parent were genotyped using the Vfaba_v2 Axiom SNP array which comprises 65,522 markers located within gene coding sequences of *V. faba* cultivar ‘Hedin 2’ (Skovbjerg et al. 2023). Genotypes were filtered by removing monomorphic markers, markers with more than 2 % missing calls, fewer than 30 homozygous calls, heterozygous parental calls, or markers for which parental replicates yielded non-matching calls other than missing calls in either replicate. Individuals were retained if they yielded more than 95 % calls. Of 173 F_2_ individuals which passed filtering, 169 were retained for mapping, with four excluded for significant segregation distortion. Linkage maps were build using R-packages ASMap v. 1.0.4 (Taylor and Butler 2017) and qtl v. 1.50 (Broman et al. 2003). The linkage map was constructed using the mstmap algorithm with p-value threshold set to 1e-9 and mapping function ‘kosambi’. The estimated genotyping error rate was 0.003. Markers were anchored to the reference genome assembly of *V. faba* ‘Hedin/2’ GCA_948472305.1 (Jayakodi et al. 2023), referred to as the reference assembly in this article, by searching flanking sequences of all markers against the genomic sequence, using blastn v. 2.15.0+ with parameters: ‘-task blastn-short’, ‘-evalue 1e-3’, ‘-qcov_hsp_perc 0.9’, ‘-subject_besthit’ and ‘-num_alignments 5’ (Altschul et al. 1990; Camacho et al. 2009).

### Phenotyping and QTL-mapping analysis

Six control varieties and 184 NV490-3 x NV512-1 derived F_3-_families (suppl. table S2) were dip-inoculated with a mixture of two isolates of *F. avenaceum* and five of *F. oxysporum* in two identical experiments conducted in separate glasshouses between March and May 2022 and between October and December 2022, respectively. Each experiment consisted of three replicates of 184 F3 families of 52 pots each, with six controls (*V. faba* cultivars ‘LG Cartouche’, ‘Fuego’, ‘Ghengis’, ‘Maris Bead’ and parental lines NV490-3 and NV512-1) replicated once within each of these blocks. Experiments were randomised using R package blocksdesign v. 4.9 (Edmondson 2020). Five seeds per pot were planted into 1 litre (L) pots with medium-grade vermiculite and watered. Pots were placed into a heated glasshouse with Sodium 10000 lux lighting for 16 h per day. *Fusarium* isolates were cultured in liquid medium made from 15 g carboxymethylcellulose sodium salt (low viscosity), 1.0 g yeast extract, 1.0 g NH_4_NO_3_, 1.0 g KH_2_PO_4_, 0.5 g MgSO_4 •_ 7 H_2_O in 1 L distilled water (Zitnick-Anderson et al. 2020) and incubated at room temperature for 21 days. Spore suspension was prepared by filtering the liquid cultures through nylon mesh (250 µM). Plant roots were dip-inoculated in spore suspension (1 × 10^5^ spores per ml) at 17 days after planting. Plants were uprooted and the roots placed in spore suspension for 15 min before repotting. Disease symptoms were scored visually as percentage of discolouration separately for stem base (foot) and roots 28 days after inoculation individually for each plant. Glasshouse temperatures averaged 22 degrees C in experiment 1 (UK springtime) and 19.2 degrees C in experiment 2 (UK autumntime). Summary statistics for foot and root scores and analyses of variance (ANOVA) were calculated in Genstat 22^th^ edition (VSNI 2022). Combined foot and root rot scores were calculated as the arithmetic mean of raw root and foot scores. Best linear unbiased predictions (BLUPs) were estimated for each of the two experiments using the H2cal function from R-package inti v. 0.6.5 (Lozano-Isla 2023), using the model:

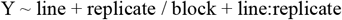

with factor ‘line’, representing the F_3_-family or cultivar, being set to ‘random’ to calculate best linear unbiased predictions (BLUPs). Factors ‘replicate’ and ‘block’ described the blocking structure. Pearson’s correlations were calculated and plotted using R package PerformanceAnalytics v. 2.0.4 (Peterson et al. 2020). The linkage map including QTL intervals was displayed using R-package LinkageMapView v. 2.1.2 (Ouellette et al. 2018).

QTLs were mapped using R-package qtl v. 1.60 (Broman et al. 2003; Broman and Sen 2009). Initial QTLs and LOD-thresholds were identified with ‘scanone’, utilising the Haley-Knott function. Logarithm of odds (LOD) thresholds were calculated at alpha 0.1, 0.05 and 0.01 in 5000 permutations. Initial QTLs, identified through ‘scanone’, were refined with ‘refineqtl’ and potential additional loci identified with ‘addqtl’ and included into the final model if they exceeded the 0.05 threshold estimated by ‘scanone’ (hk). Interactions between multiple QTLs were identified with ‘addint’ and the final QTL-model evaluated with ‘fitqtl’. Flanking markers were identified using ‘lodint’ with interval boundaries set to 1.5 x LOD either side of the peak marker. Phenotype x genotype plots were generated for each peak marker using ‘plotPXG’. Genes within the intervals were functionally annotated using InterProScan-5.67-99.0. Enrichment for GO-terms was calculated using the AgriGO v. 2 (Du et al. 2010; Tian et al. 2017) webserver (http://systemsbiology.cau.edu.cn/agriGOv2/index.php), using the GO-terms generated for the reference protein by InterProScan set for ‘Hedin/2’ as reference. Enrichment was tested using the Fisher test combined with FDR correction of Yekutieli (Benjamini and Yekutieli 2001) with a p-value threshold of 0.05. The full set of GO-terms was selected.

### Synteny with QTLs associated with disease resistance in other species

Flanking- or primer sequences of Illumina Infinium® II 15k pea array markers (Tayeh et al. 2015; Wu et al. 2022), SSR-markers (Loridon et al. 2005; Bordat et al. 2011; Mc Phee et al. 2012; Coyne et al. 2015) and KASP (Kompetitive Allele Specific amplification)-markers (Duarte et al. 2014), linked to resistance to foot rot in pea, were anchored to the faba bean reference assembly as described above for anchoring the *V. faba* Axiom-chip markers. For SNPs linked to foot rot resistance in lentil (Heineck et al. 2022), common bean (Paulino et al. 2021) and chick pea (Alsamman et al. 2024), 1 kilobase (kb) of flanking sequence was retrieved from the respective reference genome assemblies for lentil (Lculinaris_718_v1.fa of *L. culinaris* ‘CDC Redberry’ (Vandenberg et al. 2006), common bean assembly Pvulgaris_442_v2.0.fa of *P. vulgaris* ‘G19883’ (JGI, 2023) and chickpea assembly Ca_v2.6.3_kabuli_ref.fa of *C. arietinum* ‘CDC Frontier’ (Edwards 2016), using samtools faidx (Li et al. 2009) before searching the sequence against the *V. faba* ‘Hedin/2’-reference assembly using blastn v. 2.15.0+ with parameters: ‘-evalue 1e-5’, ‘-subject_besthit’ and ‘-num_alignments 5’ (Altschul et al. 1990; Camacho et al. 2009).

### Annotation of putative defensin-like genes in the *V. faba* reference assembly

Putative defensins were identified by mapping publicly available defensin protein sequences, retrieved from the NCBI protein database (https://www.ncbi.nlm.nih.gov/protein/, accessed 16 June 2024) to the reference genome using miniprot v. 0.12-r237 (Li 2023). Complete annotations of putative matching genes were generated using AGAT v. 1.4.0 scripts (Dainat et al. 2024) ‘agat_sp_add_start_and_stop.pl’ and ‘agat_sp_filter_incomplete_gene_coding_models.pl’. For identification of possibly mis-annotated short antimicrobial proteins (AMPs), RNAs identified in the reference annotation as ‘lnc_RNA’ (long non-coding RNA) were converted into messenger-RNA (mRNA)-type-annotations. Full annotations of putative protein-coding transcripts, including exons, introns, coding sequences (CDSs), stop-and start-codons were recovered and converted into GFF3-format using AGAT scripts ‘agat_sp_fix_cds_phases.pl’, ‘agat_sp_add_start_and_stop.pl’ and ‘agat_sp_filter_incomplete_gene_coding_models.pl’. Protein sequences were extracted from these additional annotations with ‘agat_sp_extract_sequences.pl’ and functionally annotated using InterProScan-5.67-99.0 (Jones et al. 2014).

Reannotating reference transcripts identified as long non-coding RNA identified 8845 putative additional coding sequences (suppl. file S2). Functional annotation of the corresponding proteins identified 26 additional putative defensins on all chromosomes except chromosome 2, and one on an unplaced scaffold (suppl. file S3). They were identified as defensin-like proteins of classes 1, 184-like and 245 (suppl. file S4).

## Results

### Phenotypic assessments of *V. faba* for *Fusarium* foot and root rot resistance

Above-ground symptoms began to appear approximately 14 days after inoculation in susceptible lines, with affected plants showing foliar chlorosis, progressing to grey, papery necrotic patches (Fig. 1a). Dark brown or black hypocotyl lesions (foot rot symptoms) appeared at the base of the stem, initially confined to the epidermis, but later penetrating deeper into the tissue. Observed severity amongst *V. faba* genotypes ranged from less than 10 % to exceeding 70 % in both experiments. Diseased roots (root rot symptoms) turned black, with blackening usually progressing upwards from the lower half of the root system (Fig. 1b). The disease scores between experiment one and experiment two were consistent for parent NV490-3, with foot and root rot symptoms averaging around 43 % in both experiments. A genotype effect, represented by factor ‘line’ was significant in both experiments, for severity of foot and root rot symptoms (*p* < 0.001, suppl. table S4). Parent NV512-1 showed much more severe symptoms in experiment one, with 54 % and 61 % for foot and root damage respectively and 39 % and 43 % in experiment two (suppl. table S3, Fig. 2a, b). Four F_3_-families, NV490-3xNV512-1_77, NV490-3xNV512-1_80, NV490-3xNV512-1_85 and NV490-3xNV512-1_30, demonstrated greatly reduced disease symptoms compared to either parent; these lines showed levels of damage between 3 % and 24 % for both traits, whilst other lines showed little hypocotyl damage but more severe root damage, e.g. NV490-3xNV512-1_137 whilst the reverse was not observed (suppl. table S3). The most susceptible families exceeded the damage seen in the NV512-1 susceptible parent e.g. NV490-3xNV512-1_4, yielding disease scores greater than 74 % for root and foot rot symptoms in both experiments. Control cultivar ‘Maris Bead’ also showed consistently low levels of disease symptoms for both tissues, with between 12 % and 17 % hypocotyl damage and 21 % and 40 % root damage in experiments one and two, respectively. The two types of symptoms were strongly correlated within each experiment (*R2* = 0.76) (Fig. 2 a, b) however, only a minor correlation (*R2 =* 0.43) between symptoms was observed between experiment one and two (Fig. 2 c, d).

**Figure 1.**
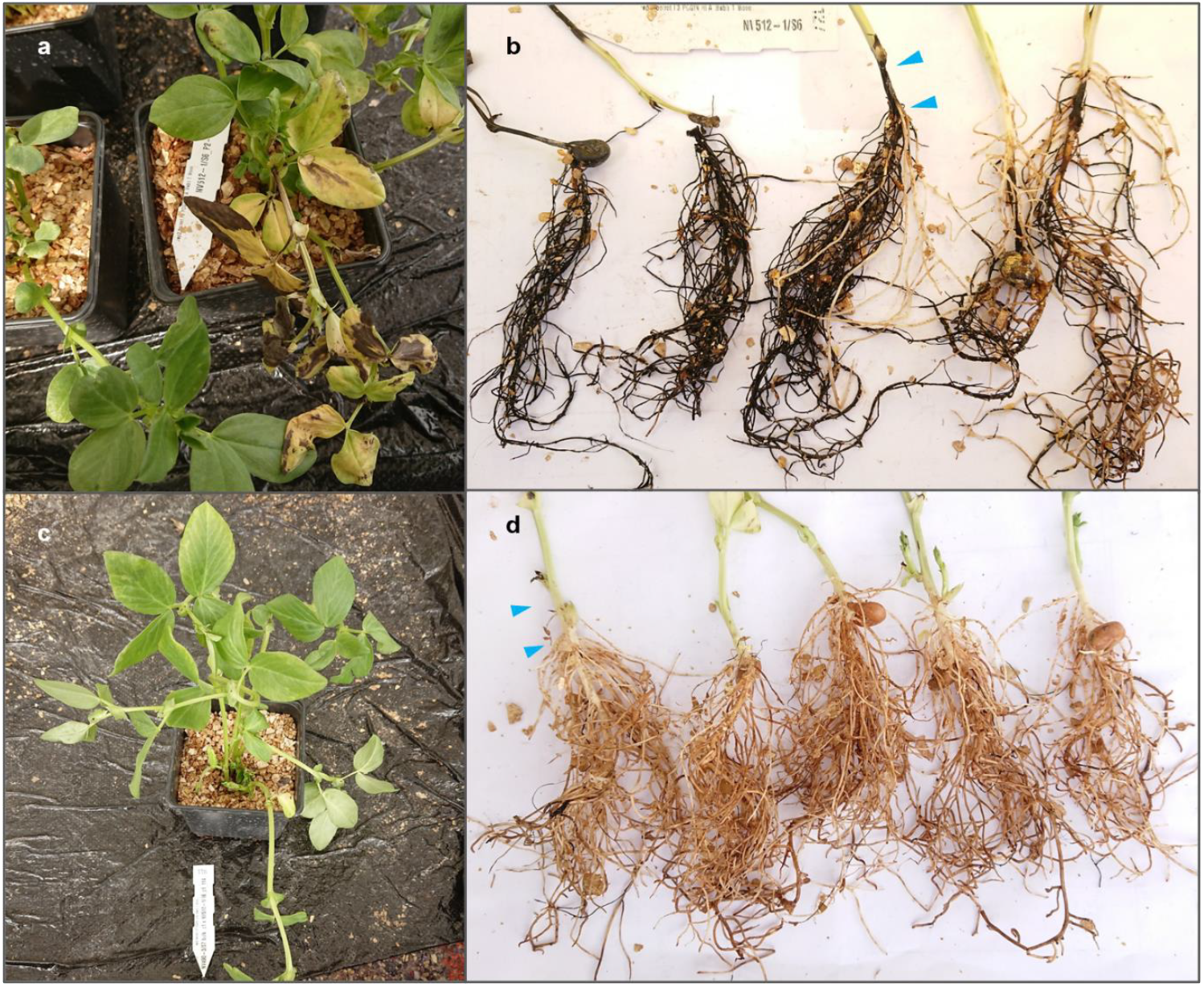
Foot and root rot symptoms in *Vicia faba*, first experiment. Above-ground symptoms in susceptible parent NV512-1 (a), blackening of stem base tissue (foot, blue arrows) and root system (b), resistant F_3_ family NV490 x NV512-114 shoots without obvious foliar symptoms (c), and largely healthy stem base and root system (d)

**Figure 2.**
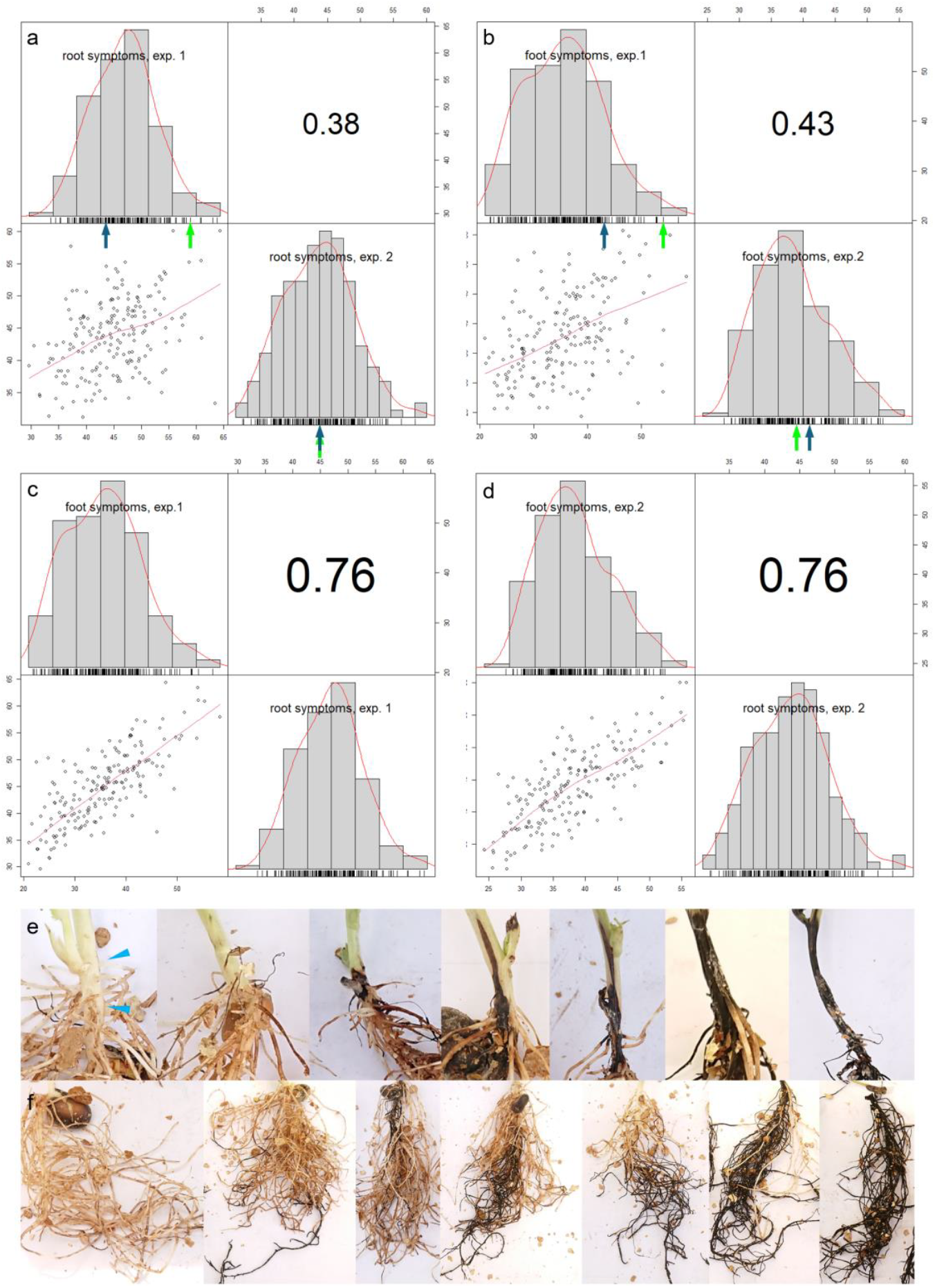
Pearson’s correlations of best linear unbiased predictors (BLUPs) between disease symptoms, comparing experiments, and foot (stem base) and root symptoms. Top row: correlations of foot (a) and root symptoms (b) between experiments, with parents NV490-3 (blue arrows) and NV512-1 (green arrows) indicated below histograms. Middle row: correlations between root (c) and foot symptoms (d) in experiment one (exp. 1) and experiment two (exp. 2). Diagonal graphs show histograms of BLUPs, name of trait or experiment. Bottom left rectangle: scatter plots, top right rectangle: correlation coefficients. P-values for all Pearson’s correlation tests < 0.001. Graph generated with R-package PerformanceAnalytics v.2.0.4 (Peterson et al. 2020). Bottom: scales of foot rot symptoms on stem base (light blue triangles) (e) and root rot symptoms (f) observed in the experiments, from left to right: 0-5 %, 6-10 %, 10-25 %, 25-50 %, 50-75 %, 75-95 %, 100 %

### Linkage map construction and QTL analysis

The final linkage map consisted of 6577 markers in seven linkage groups (LG) and has a length of 2145.34 cM. Linkage group indices 1 to 6 correspond to the six chromosomes of *V. faba* to which their constituent markers were predominantly mapped Chromosome 5 is represented by two linkage groups, LG5.1 and LG5.2 (suppl. table S2, Fig. 3). This was due to a region of segregation distortion corresponding to 61.2 Mb to 1140.1 Mb on chromosome 5 in the reference assembly (suppl. table S5, *χ*^2^ test p < 0.05). The region between 638.4 Mb and 949.4 Mb of the chromosome in the reference assembly was not covered by any markers suitable for mapping within this population. A progressive reduction in homozygous NV512-1 allele (BB) calls and concomitant increase in the homozygous NV490-3 allele (AA) calls could be observed with a complete absence of BB-calls in a region spanning approximately 310 Mb, corresponding to 638.4 Mb and 949.4 Mb on the reference assembly, before normalizing (suppl. Fig. S1, suppl. table S5).

**Figure 3.**
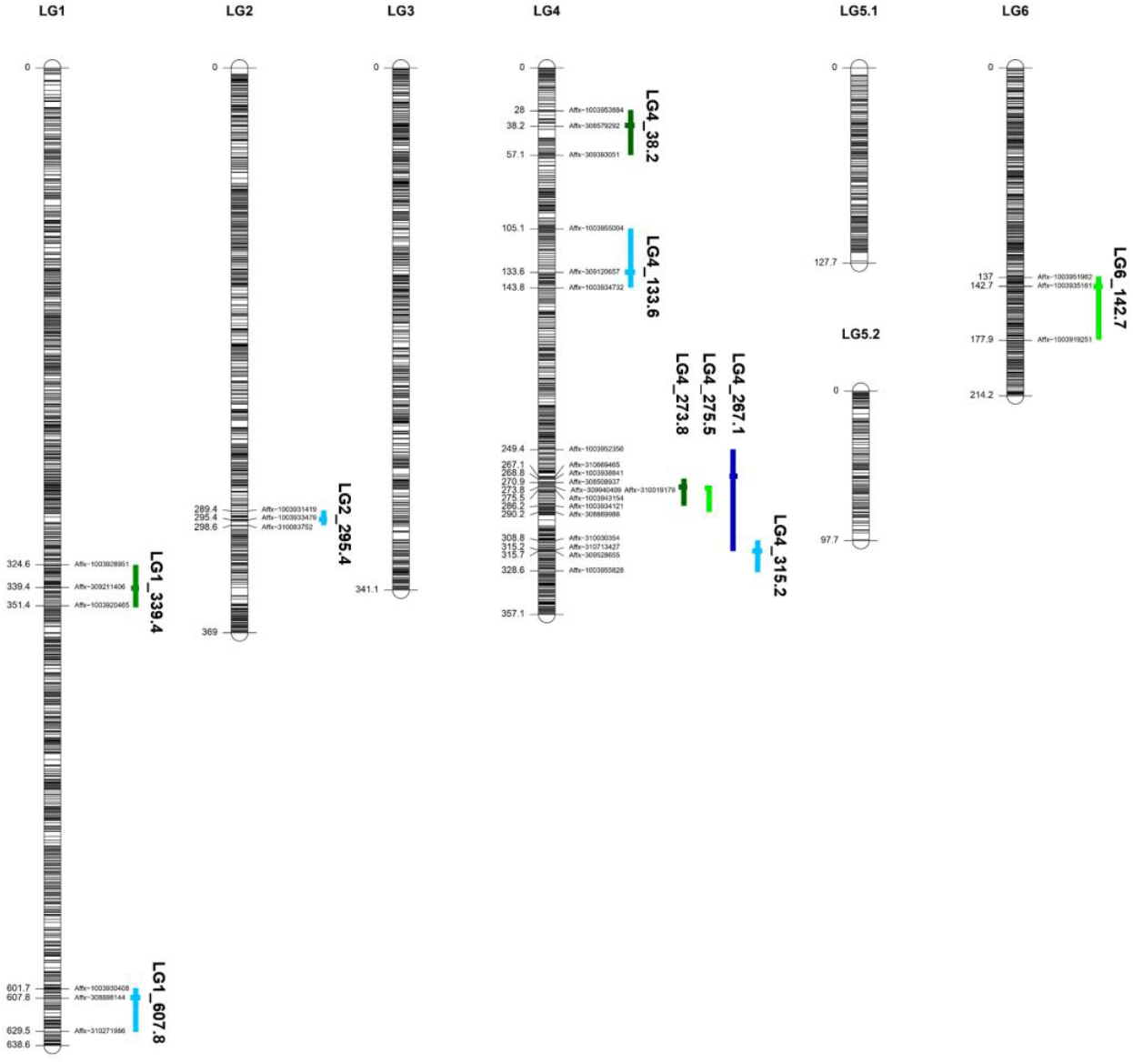
Linkage map of NV490 x NV512, with QTLs for foot rot symptoms (light green experiment one, dark green experiment two) root rot symptoms (light blue experiment one, dark blue experiment two) and displayed to the right of their respective loci. Names and positions in cM of flanking and peak markers are shown for each QTL. Map plotted with R-package LinkageMapView v. 2.1.2 (Ouellette et al. 2018).

The closest mapped markers flanking the distorted region of chromosome 5 were Affx-1003947844 at 118.919 cM of LG 5.1, corresponding to 638.4 Mb, and Affx-1003916494 at 97.690 cM of LG5.2, located at 949.3 Mb. The region between these markers contains 1102 annotated genes. Markers Affx-1003943804, Affx-1003927953 and Affx-308839474, located at 843.56 Mb, 844.48 Mb and 848.00 Mb in the centre of the distorted region returned no homozygous calls for the NV512-1 allele within any of the progeny. A further eleven markers, between Affx-309772554 (804.88 Mb) and Affx-1003919956 (848.59 Mb), flanking those three, only returned one homozygous NV512-1 allele call each. The region covered by these fourteen markers contains 131 genes encoding 114 putative proteins and 23 non-coding RNAs (suppl. table S6

### Identification of QTLs associated with *Fusarium* foot and root rot resistance

Two QTLs associated with resistance towards foot rot were detected on LG4 (LG4_275.5) and LG6 (LG6_142.7) respectively, in experiment one. These accounted for 19.8 % and 8.8 % of the phenotypic variation explained (PVE), respectively. However, only the locus on LG4 was detected again in the second experiment, with QTL LG4_273.8 cM, accounting for 12.7 % of the PVE. Two additional loci, LG1_339.4, andLG4_38.2 were identified in experiment two, accounting for 9.0 % and 8.7 % PVE respectively.

For the root rot phenotype, two independent QTLs, LG4_133.6 and LG4_ 315.4 were observed to accounted for 13.2 % and 16.2 % of the PVE, respectively, in experiment one. Two additional loci LG2_295.4 andLG1_607.8, were observed and accounting for 10.0 % and 8.0 % PVE, respectively. In experiment two, only a single QTL, LG4_ 267.1, was detected, accounting for 10.6 % PVE.

All the QTLs detected in the study were inherited from the resistant parent (suppl. figs. S1, S2), except for LG2_295.4, associated with the root rot phenotype, for which NV512-1 was the donor (suppl. fig. S2 b).

QTLs LG4_273.8, LG4_275.5 (foot rot) and LG4_267.1 (root rot) may correspond to the same underlying locus, with the remaining QTLs on LG4 corresponding to separate loci.

For the combined foot rot scores QTLs were identified on LG4 in both experiments at LG4_285.3 in experiment1 and at LG4_273.8, accounting for 16.1 % and 11.6 % PVE respectively, with intervals overlapping each other and the cluster of 3 QTLs on LG4 between 267 cM and 275 cM associated with foot- and root rot in both experiments. Additional QTLs LG6_141.0 (exp. one, 8.3 % PVE) and LG1_341.8 (exp. two, 8.7 % PVE) likely correspond to QTLs LG6_142.7 and LG1_339.4, associated with foot rot symptoms (suppl. table S6). Combining the foot and root rot scores led to a reduced resolution for all observed QTLs, therefore, the two traits were reported separately.

**Table 1.**
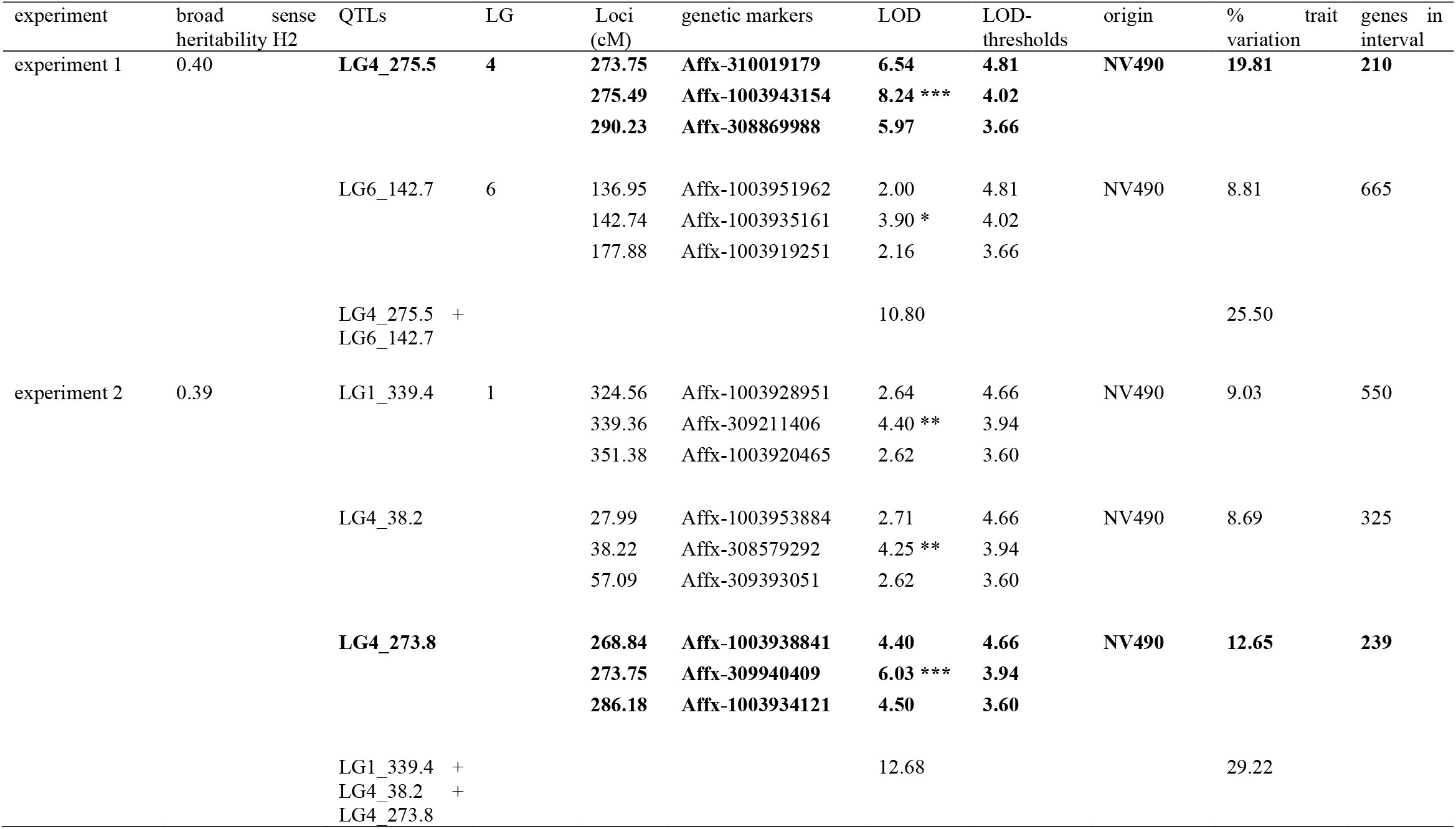
QTLs detected for foot rot symptoms in experiment one and experiment two. LOD-thresholds at α 0.01, 0.05, 0.1, calculated with R-qtl function scanone, Haley-Knott, 5000 permutations. Broad-sense heritability H2 after Cullis et al. (1996), calculated with H2cal function of R-package inti (Lozano-Isla 2023). The resistant genotypes for all QTLs are derived from parent NV490. Text marked in bold indicates QTL consistent across both experiments.

**Table 2.**
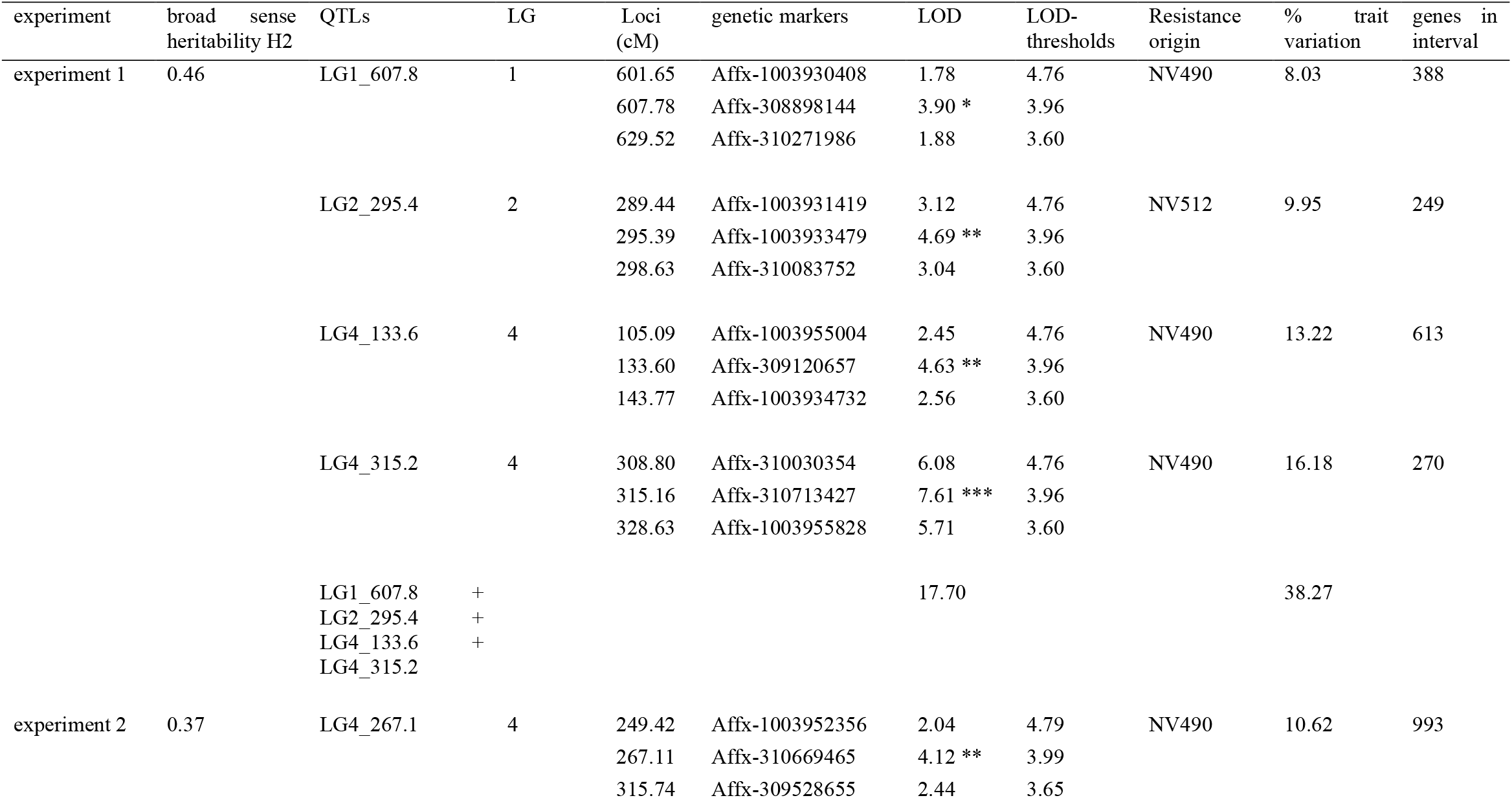
QTLs detected for root rot symptoms in experiment one and experiment two. LOD-thresholds at α 0.01, 0.05, 0.1, calculated with R-qtl function scanone, Haley-Knott, 5000 permutations. Broad-sense heritability H2 after Cullis et al. (1996), calculated with H2cal function of R-package inti (Lozano-Isla 2023). The resistant genotypes for all QTLs except for QTL LG2_295.4 are derived from parent NV490.

### Candidate genes identification and GO-term enrichment

QTLs LG4_273.38 and LG4_275.5 were enriched for GO-terms relating to regulation of cell cycle, DNA-damage, photosynthesis and response to stress (suppl. table S8), whilst QTL LG4_267.1 was enriched for functions relating to enzyme regulation and inhibition. Several genes putatively associated with defence against pathogens were located within the QTL intervals, e.g. one gene annotated as innate immune response and plant defence kinase, and a cluster of four pathogenesis-related proteins of the *Bet v 1* family. Two clusters of genes annotated as exostosin */* heparan sulphate glycosyltransferase-related and glycosyl hydrolase family 79 */* heparanase-related were found to be adjacent to the peaks of LG4_273.8 and LG4_275.5.

Similarly, QTL LG4_133.6 contained a cluster of seven defensin-like protein 1 / Knottin, scorpion toxin-like superfamily / gamma-thionin superfamily genes, six of which are homologous to *Ec-AMP-D1, Ec-AMP-D2, SD2*-like or *MtDef4*.*2*-like defensins, found in other legumes (suppl. table S9). Four TIR-NBARC-LRR (TNL)-type genes (one TMV resistance N-like, one *RUN1*-like and two *RPV1*-like resistance genes) and two chitinases are also located under the QTL. LG4_133.6 was enriched for 66 GO-terms, e.g. nucleotide synthesis, transmembrane transport, oxidoreductase activity and carbohydrate-binding processes, and cellular components plant cell wall and proton-transporting ATP-complex. LG4_33.8 was enriched for GO-terms relating to pectin metabolism, gene expression and transcription and lipid-binding functions.

QTL LG6_142.7 contained three defensins homologous to *MtDef4*.*6*, Defensin 1-like and fabatin, and a cluster of nine thaumatin-family secreted proteins and one *Gnk2* / salt stress antifungal protein. The region is enriched for GO-terms relating to transcription-related processes and transcription factors.

QTL LG1_339.4 overlays eight small non-specific lipid transporters of which seven were clustered near its peak. It also contained three TNLs annotated as TMV-resistance protein N-like. The region was enriched for GO-terms relating to cellular component plasma membrane. The second QTL on LG1, LG1_607.8, overlayed one small secreted antifungal protein annotated as pollen protein *Ole e 1*-like, a large cluster of cysteine protease family C1-related, two CNLs and one *Gnk2 /* salt stress antifungal protein, a cluster of fourteen *PR1*-like secreted proteins and two OS01G0750732 proteins annotated as innate immune response and plant defence kinase. This locus is not significantly enriched for any GO-terms.

The only QTL derived from NV512-1, LG2_295.4, was enriched for 59 GO-terms relating to root and shoot system development and several metabolic processes. It contained one chitinase-1 gene and several plant LRR receptor-like serine/threonine-protein kinases.

## Discussion

The aim of our study was the identification of loci conferring broad-spectrum resistance to a combination of *Fusarium* species causing foot and root rot in the UK. No QTLs associated with resistance to foot and root rot have previously been reported in *V. faba*. We describe the first QTLs associated with partial broad-spectrum resistance to foot and root rot caused by *Fusarium* spp. in a bi-parental mapping population of *V. faba*. In several other legumes QTLs conferring resistance to Fusarium wilt or foot and root rot have previously been identified, e.g. in pea (Grajal-Martìn and Muehlbauer 2002; Mc Phee et al. 2012; Coyne et al. 2015, 2019; Wu et al. 2022), chickpea (Jendoubi et al. 2016; Garg et al. 2018) and common bean (Nakedde et al. 2016; Wang et al. 2018; Paulino et al. 2021). One study had identified potential sources of partial resistance in *V. faba* to wilt caused by *F. oxysporum*, but no QTLs were identified (Mahmoud and Abd El-Fatah 2020). In our experiments, a wide range of foot rot and root rot severity were observed among the progeny with scores exhibiting near-normal distributions for both phenotypes. Levels of resistance or susceptibility in some progeny exceeded the levels observed in the parental lines, pointing to an involvement of multiple, small-effect loci, originating from both parents, as commonly observed for quantitative resistance to necrotrophic pathogens with multiple host species (Mengiste 2012). Most F_3_-progeny and the resistant parent NV490-3 showed between 35 % and 45 % root or foot blackening in both experiments with some progeny either showing less than 20 % disease symptoms or exceeding the disease scores of the susceptible parent NV512-1. Four F3 families with promising levels of resistance to foot and root rot were identified in this study. Despite having been conducted under controlled environmental conditions, substantial variation between the two experiments was observed, which likely affected the reproducibility and precision of mapping small-effect QTLs. Differences in maximum temperatures were a likely cause for some of the variation observed between the experiments. Experiment one was exposed to higher day time glasshouse temperatures in May 2022 compared to the second experiment, which concluded in the autumn. *Fusarium* species involved in foot and root rot of legumes, including pea (Esmaeili Taheri et al. 2017) and faba bean (Šišić et al. 2022) have been found to favour different environmental conditions, and this could have played a role in the experimental variation described herein. Whilst efforts were made to equilibrate the quantity of spores from each isolate used for screening, the composition of the inoculum, specifically the ratio of *F. avenaceum* to *F. oxysporum* spores, and segregation within individual plants comprising F3 families in genes contributing to resistance could also have affected the level of disease symptoms that were observed. Subsequent independent testing of *F. avenaceum* and *F. oxysporum* on selected lines from the NV 490 × 512 mapping population, demonstrated that levels of disease were comparable to those observed in the experiments reported here, supporting the case that QTL identified in the study confer resistance against both species. Studies conducted with single *Fusarium* species and within more environmental conditions would be needed to verify and refine the identified QTLs and possibly identify additional loci. Also, repeating the experiments in recombinant inbred lines derived from the cross would likely facilitate fine mapping of loci involved in foot rot resistance and help to mitigate symptomatic variation due to genetic heterogeneity.

Multiple small-effect QTLs associated with resistance to foot and root rot were observed to account individually for between 8 % and 20 % in trait variation. The locus with the greatest effect associated with resistance to foot and root rot was identified in both experiments and is covered by QTLs LG4_275.5, LG4_273.8 (foot rot, experiments one and two) and LG4_267.1 (root damage, exp. one) accounting for between 12 % and 20 % of phenotypic variation. LG4_315.2 (root damage, exp. two) overlapped with LG4_267.1, but not with either of the two QTLs associated with levels of foot rot and may therefore correspond to a separate locus on LG4. The region covered by QTLs LG4_273.8 and LG4_275.5 contained 239 annotated genes. Two clusters of genes encoding proteins annotated as possessing transmembrane domains and functions “exostosin, heparan sulphate glycosyltransferase-related” and “Glycosyl hydrolase (GH) family 79, heparanase-related” respectively, are located between the two peak markers. In animals, heparanases are known to be involved in the remodelling of cell membranes and involved in signalling pathways in response to inflammation through damage associate molecular patterns (DAMPs) (Masola et al. 2018), a mechanism also known to be involved in the PTI-defence response (Pattern Triggered Immunity) of plants to pathogens (Hou et al. 2019). To date their function in plants has not been characterised yet.

All other QTLs observed in the study were only detected in a single experiment. No QTLs associated either with foot rot or root rot severity covered a common locus, suggesting that the genes residing in them may correspond to tissue specific resistance. They accounted for between 8 % and 13 % of observed trait variation. Although root and foot symptoms were strongly correlated in both experiments, some progeny showed differential scores for root and foot rot, suggesting that multiple genes are contributing to broad-spectrum resistance observed in this population, some tissue specific, some not. Most of these QTLs contained several classes of genes linked to broad-spectrum resistance to pathogens, some of which have been found to be involved in resistance to *Fusarium* spp. in other legume species.

Two of the QTLs identified in experiment one, LG6_142.7 (foot rot) and LG4_133.6 (root rot), could be successfully anchored to loci previously associated with resistance to necrotrophic pathogens in faba bean and pea. LG4_133.6 corresponds to a region on *V. faba* chromosome 4 syntenous to chromosome 4 of *P. sativum* ‘Cameor’ which contains QTLs *Fg-Ps4*.*1* and *Fg-Ps4*.*2* associated with severity of foot rot in pea (Wu et al. 2022). QTL LG6_142.7 overlaps a region on the *V. faba* ‘Hedin/2’ reference genome to which QTLs LG6_121.0 and LG6_169.6, associated with resistance to chocolate spot, caused by *Botrytis fabae*, have been mapped (Webb et al. 2024). Loci residing in these regions have been associated with partial resistance to chocolate spot in faba bean cultivar ‘Maris Bead’, indicating that they may be associated with broad-spectrum resistance to necrotrophic pathogens, or that genes regulating resistance to necrotrophic pathogens are situated in similar locations in the faba bean genome. The remaining QTLs, including the locus on LG4 found in both experiments, could not be anchored to any of the previously described QTLs associated with resistance to foot rot in other legumes, indicating they likely represent novel sources of *Fusarium* resistance.

All the QTLs identified in a single experiment within study overlay clusters of small secreted antimicrobial proteins (AMPs) which have been associated with plant defence against necrotrophic pathogens, including several *Fusarium* species (Goyal and Mattoo 2014). QTLs LG4_133.6 and LG6_142.7 overlap clusters defensin-1 type / knottin-like secreted proteins which have been identified as candidate genes involved in defence against *F. oxysporum* f. sp. *pisi* in pea (Coyne et al. 2015). Defensins are small cysteine-rich proteins involved in innate immunity across all forms of cellular organisms (Thevissen et al. 2000). In plants, defensins function as part of the innate broad-spectrum defence and have been found to display anti-fungal activity and contribute to resistance to a diverse range of biotrophic and necrotrophic plant pathogens, including *Fusarium* spp. (Thevissen et al. 2000; Thomma et al. 2002; Khan et al. 2014; Goyal and Mattoo 2014; Sher Khan et al. 2019; Slezina et al. 2021).

Mapping of known plant defensin sequences to the defensins annotated on chromosome 4 under LG4_133.6 in this study showed highest sequence homology to *MtDef4*.*2* and *SD2*, previously found to inhibit *F. culmorum* (Sotchenkov et al. 2005), *F. oxysporum* and *F. solani* (Mirakhorli et al. 2019), and *F. graminearum, F. proliferatum* and *F. verticilloides* in vitro (Kaur et al. 2012). The three defensin-1 type / knottin like secreted proteins under LG6_142.7 are homologous to *MtDef4*.*6* and fabatin, a type of *V. faba* defensin found to inhibit the growth of bacteria (Zhang and Lewis 1997). Several other classes of genes involved with PTI were found underneath these two QTLs, including salt stress response/antifungal / cysteine-rich receptor-like protein kinase 3, involved in PTI against *Ustilago maydis* (Ma et al. 2023) and *Gnk2*-like secreted proteins, shown to inhibit the growth of *F. oxysporum* and other pathogenic fungi (Miyakawa et al. 2014). Also, one cluster of secreted neprosin activation peptide/glutamic endopeptidase proteins is located next to the peak of LG4_133.6. In pitcher plants (*Nepeta* spp.) neprosins function as proteases secreted into the pitcher fluid to digest the plants’ insect prey, but their function in non-carnivorous plants is currently unknown, despite them having been found in more than 100 plant species so far (Ting et al. 2022).

QTLs LG1_339.4 (foot rot, experiment two) and LG2_295.4 (root, experiment one) overlay clusters of non-specific lipid transfer (NLP) proteins, a class of anti-microbial proteins associated with resistance to root rot (*Cochliobolus sativus*) and head blight (*F. graminearum*) in wheat (Zhu et al. 2012). LG1_607.8 (root, experiment two) overlays a cluster of pathogenesis related-1 like (*PR1*-like) and *Gnk2*-like genes. *PR1*-like are thought to be involved in defence signalling in response to pathogen attack in plants (Breen et al. 2017) and have been shown be expressed in tomato in response to infection with *F. oxysporum* (Slezina et al. 2021). A single QTL LG2_295.4, was attributed to the susceptible NV512-1 parent and was enriched for GO terms associated relating to root development, chitinase functions and several plant LRR receptor-like serine/threonine-protein kinases; this locus was likely to account for the transgressive segregation observed in the most resistant progeny. Whilst synteny was observed with QTLs mapped in other legumes and the classes of genes found within their intervals suggest that these singular QTLs are indeed linked to resistance to foot and root rot, they need to be confirmed in additional experiments to evaluate their contribution to resistance to foot and root rot in faba bean.

Despite generating a densely covered linkage map for the NV490-3 x NV512-1 population, it was not possible to resolve an area of segregation distortion (SD) on LG 5, resulting in splitting the markers mapping to chromosome 5 into two linkage groups, named LG5.1 and LG5.2; no markers could be assigned to these LGs within a region of 297 Mb due to the severity of the SD. When comparing the allele frequencies, there was a significant and progressive reduction in NV512-1 calls in the F_2_ lines around the region, and a significant increase in NV490-3 calls. For three markers, located between 843.6 Mb and 848.0 Mb on chromosome 5, no calls homozygous for the NV512-1 allele were observed among the F_2_, indicating that one or multiple genes residing within this interval, were potentially contributing to incompatibility in combination with other yet unidentified loci from NV490-3. Nevertheless, as no significant QTLs associated with *Fusarium* resistance were identified on linkage groups LG5.1 or LG5.2, this anomaly did not hamper the investigation.

### Summary

This is the first study describing mapping QTLs linked to partial, broad-spectrum resistance against *Fusarium* spp. in *V. faba*, demonstrating that multiple QTLs, each contributing moderate effects, are likely to be involved in regulating this trait. One QTL associated with resistance to foot rot on LG4 was found in both experiments and overlapped with one additional QTL associated with root rot. Foot and root rot symptoms were strongly correlated, however, most QTLs associated with either foot or root rot severity mapped to separate loci, in different locations, suggesting that at least some loci may be linked to tissue-specific resistance.

The QTLs identified in this study correspond to regions containing classes of genes linked to broad-spectrum immunity to pathogens, including defensins and other small secreted anti-microbial peptides identified as candidates against *Fusarium* root rot in other legumes. Further work is needed to confirm and refine the loci found in this study, identify additional sources of resistance among accessions of *V. faba* and conduct further mapping studies involving single species of *Fusarium* involved in foot and root rot of faba bean. This will facilitate the selection and pyramiding of small-effect QTLs into elite new cultivars with high-level partial resistance against *Fusarium* species affecting faba bean.

## Data availability

The linkage map and sequence information for all mapped markers are freely available on www.viciatoolbox.org. Supplementary files are available at Zenodo: https://zenodo.org/records/14906146.

## Contributions

Jane Thomas, Anne Webb, and Thomas Wood conceived the study and designed the experiments. Anne Webb Thomas Wood, and Jane Thomas wrote and edited the manuscript. Jane Thomas obtained the funding. Anne Webb, Huw Davis, Simon McAdam and Krystyna Gostkiewicz conducted the experiments. Anne Webb prepared the DNA for genotyping, constructed the linkage map, conducted QTL mapping, conducted the bioinformatics and statistical analyses. Deepti Angra and Donal O’Sullivan developed the ‘Vfaba_v2’ Axiom SNP array and conducted the genotyping and allele calling of the samples. Martina Capozzi conducted the isolate species determination. All authors co-edited and contributed the manuscript.

## Acknowledgements

Many thanks to Tally Wright for advising on the statistical analysis and to Kostya Kanyuka for reviewing the manuscript

